## Supplementary Figures for "ISOTOPE: ISOform-guided prediction of epiTOPEs in cancer"

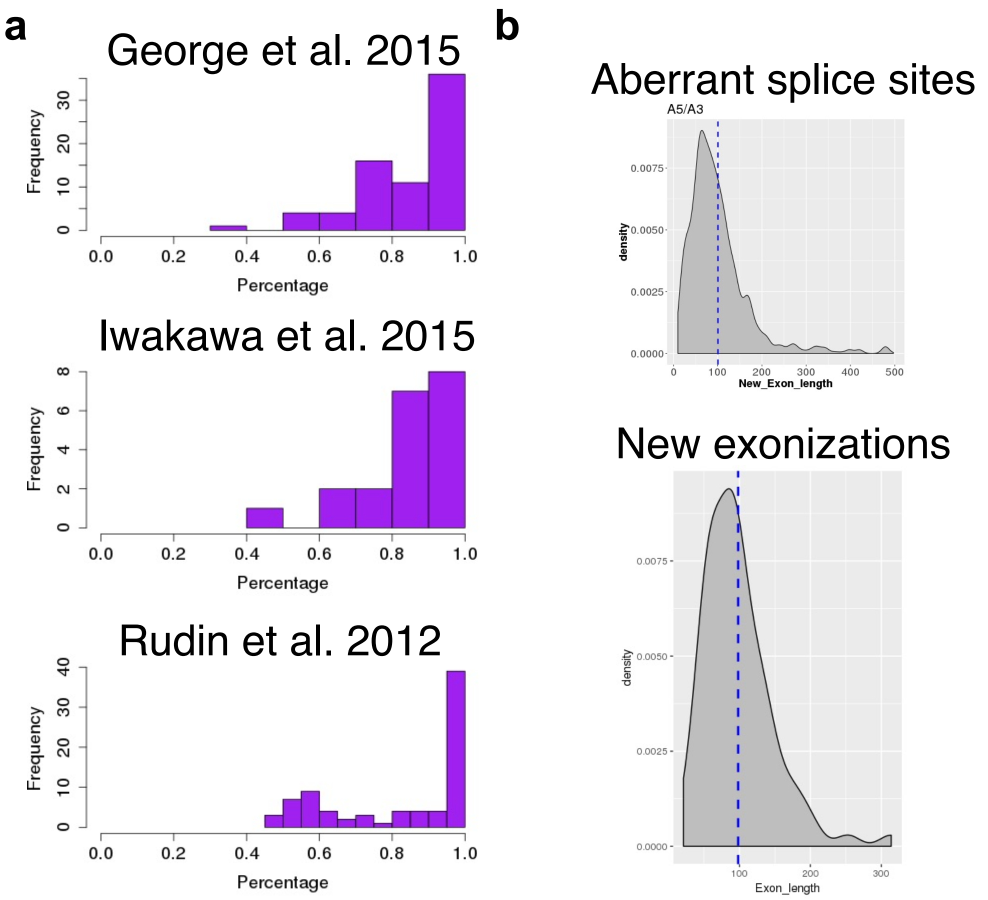


**Supp. Figure 1. Properties of the SCLC samples. (a)** Purity analysis of the small cell lung cancer (SCLC) samples calculated with ESTIMATE (Yoshihara et al. 2013). For each one of the three cohorts used for this study, we give the distribution of tumor purity values (between 0 and 1). **(b)** Length distributions of the new exons produced as a consequence of aberrant splice sites (upper panel) or new exonizations (lower panel). The lengths follow extreme value distributions with mean values of 100, similarly to known exons.

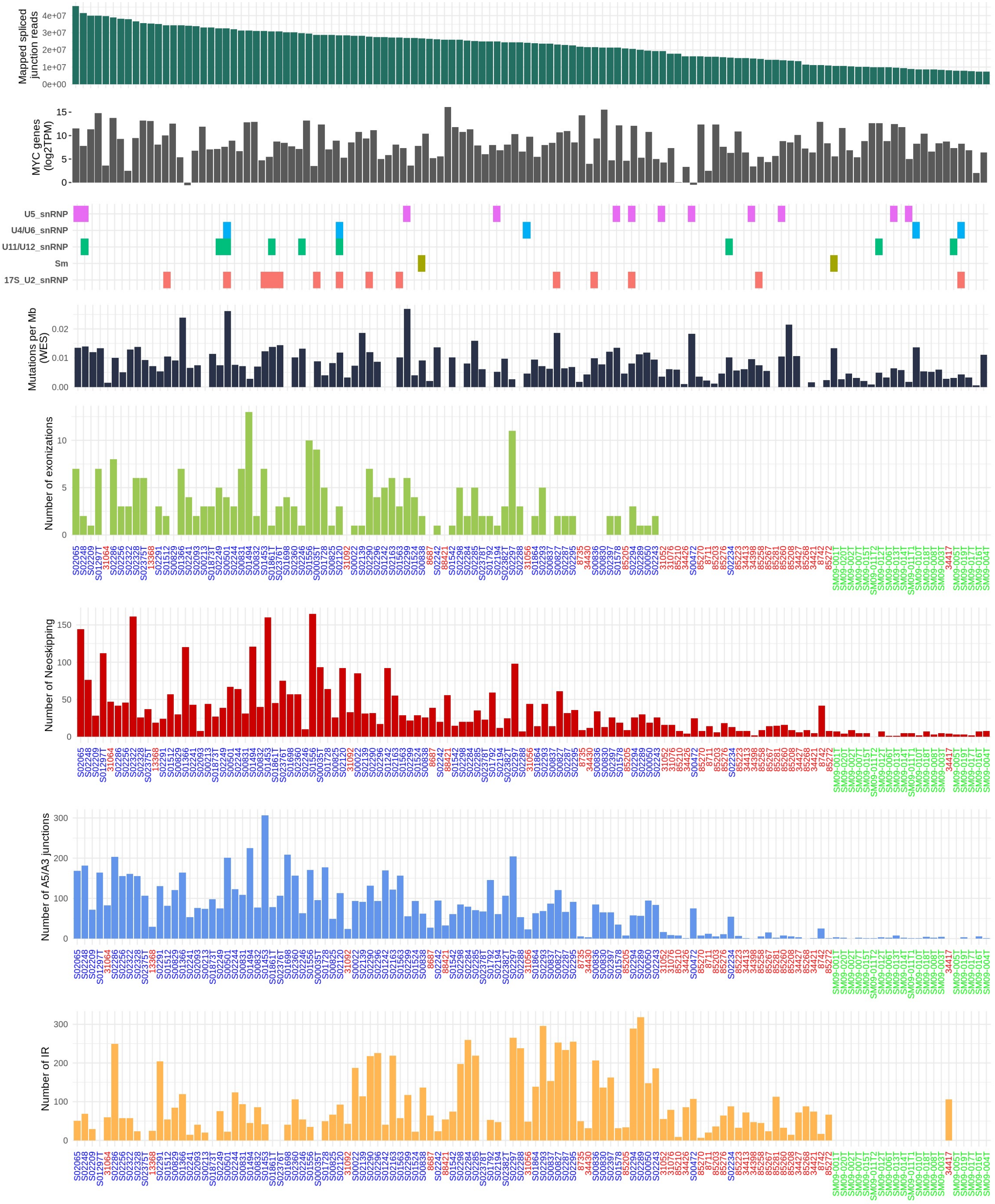


**Supp. Figure 2. SCLC-specific splicing alterations.** For each of the SCLC samples analysed (blue from George et al. 2015, red from Rudin et al. 2012, green from Iwakawa et al. 2015), we plot, from top to bottom, the number of mapped spliced reads, the expression of the MYC genes (known to be amplified or overexpressed in SCLC and to drive splicing alterations), mutations on core spliceosome factors, tumor mutation burden and the number of the different event types detected by ISOTOPE.

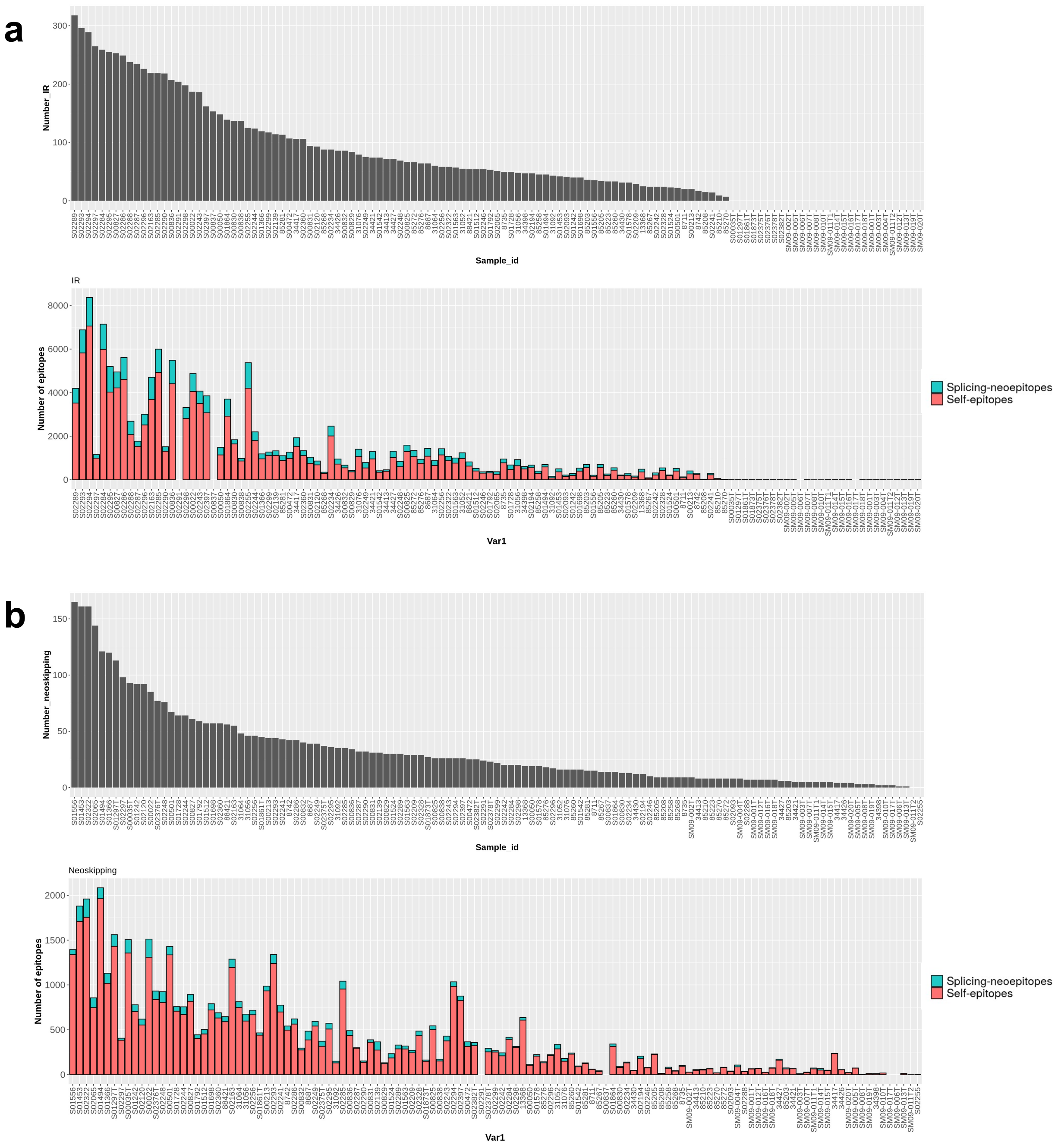


**Supp. Figure 3. Splicing-derived epitopes and splicing-affected self-epitopes in SCLC patients.** **(a)** Upper panel: Number of intron retentions per SCLC sample that impact the open reading frame. Lower panel: Number of candidate MHC-I binders per sample that are created (blue), i.e., splicing-derived neoepitopes, or potentially removed from the ORF by the splicing alteration (red) through exonizations. **(b)** Same as in (a) but for neoskipping events.

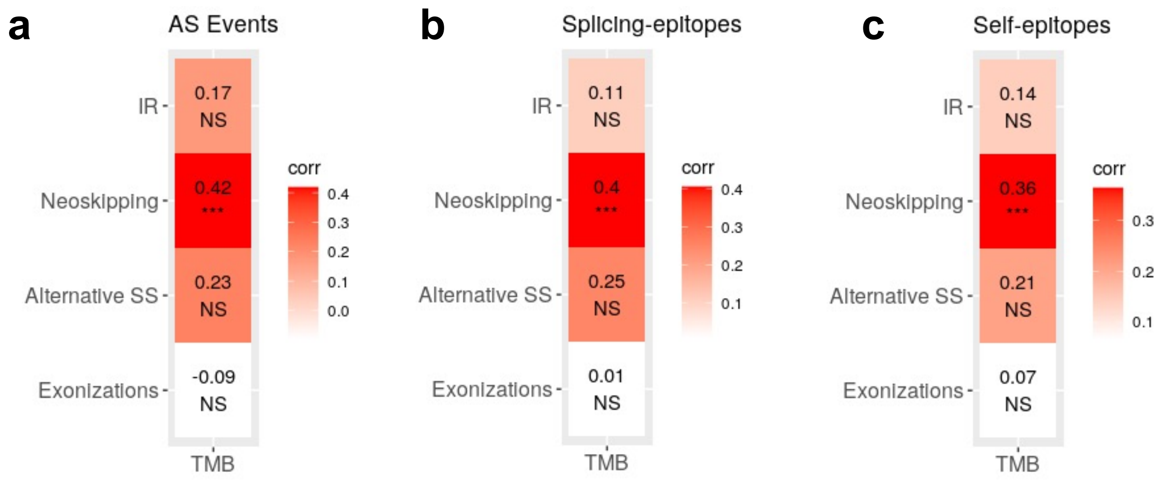


**Supp. Figure 4. Correlation of events and neoepitopes with the tumor mutation burden. (a)** Correlations between the number of splicing alterations detected and the tumor mutation burden (TMB) for all the SCLC patients (Supp. Fig. 2), separated by splicing alteration type. Although across all the events types the correlation is low (Spearman $\rho = 0.182$), separately there was a statistically significant correlation for Neoskipping events ($\rho = 0.42$). We show the same correlations separating splicing-derived neoepitopes **(b)** and splicing-affected self-epitopes. **(c).** Although neoskipping events showed significant association, there was an overall low correlation across all the event types between the TMB and the splicing-neoepitopes ($\rho = 0.194$) and self-epitopes ($\rho = 0.196$).

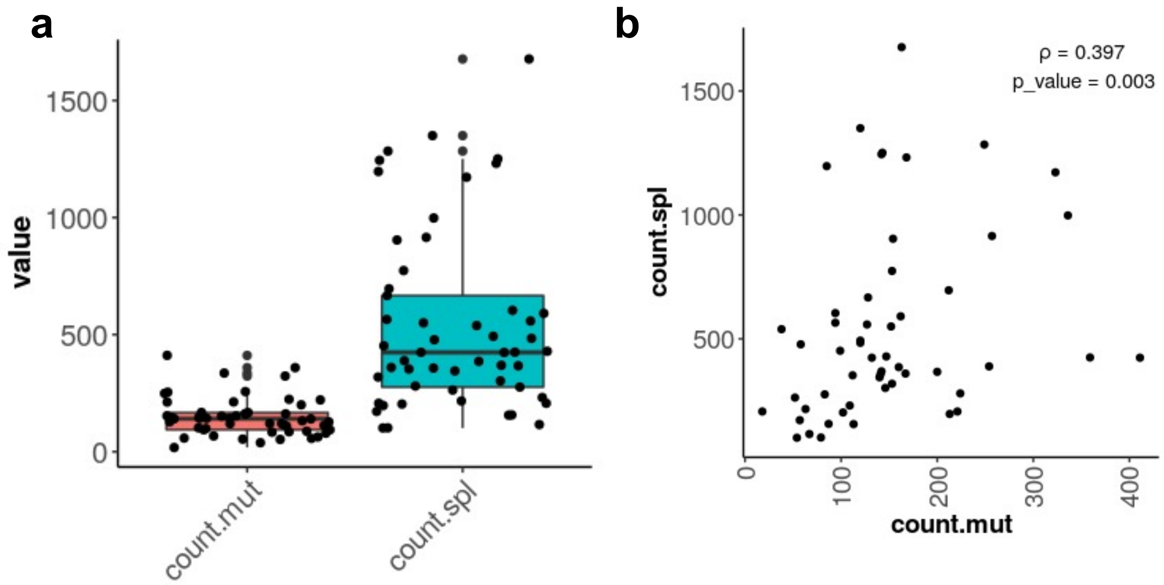


**Supp. Figure 5. Comparison between splicing-derived neoepitopes and neoepitopes derived from somatic mutations. (a)** Comparison of the number of neoepitopes derived from somatic mutations (red) and tumor-specific splicing-derived neoepitopes (blue) in the SCLC patient cohorts from (Peifer et al. 2012) and (George et al. 2015). **(b)** For each patient from the same cohorts, we give the number of neoepitopes derived from somatic mutations (x axis) and the number of neoepitopes derived from tumor-specific splicing alterations (y axis). Mutation-derived epitopes were calculated with pVACtools (Hundal et al. 2020). The identification of splicing-derived neoepitopes was carried out with ISOTOPE as described in the manuscript. The candidate epitopes were calculated in both cases using the same tools with the same parameters: NetMHC and NetMHCPan, using the hg19 reference, testing peptides with amino acid length from 8 to 11, and selecting candidates with binding affinities less or equal than 500nM. There were no overlaps between candidates generated by both methods.

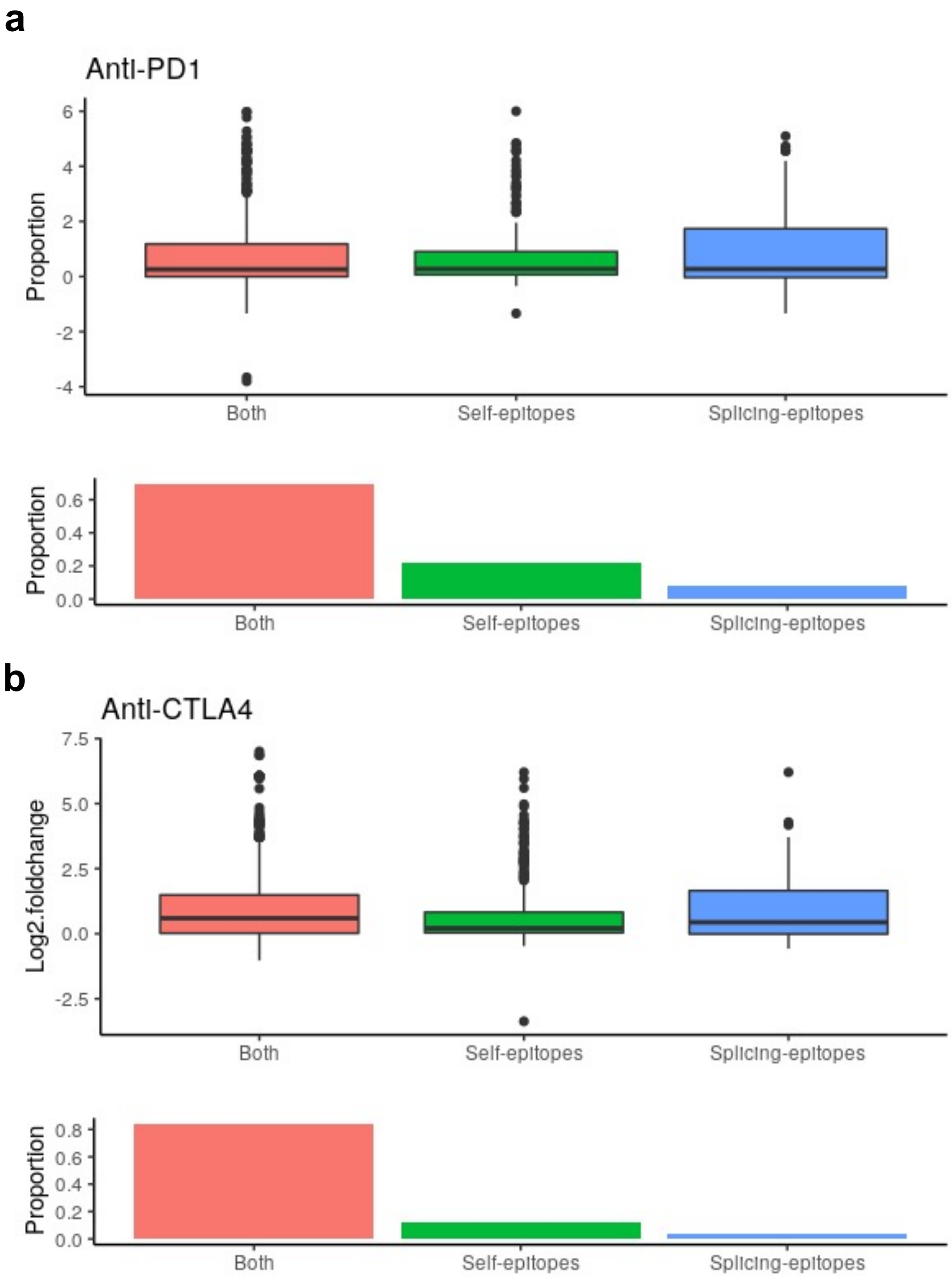


**Supp. Figure 6. Length differences between the wild type (WT) open reading frame (ORF) and the splicing altered ORF.** We show the distributions of the length ratios between WT ORF and the ORF affected by the splicing alteration for the anti-PD1 **(a)** and the anti-CTLA4 **(b)** cohort. The ratios are plotted in log2 scale, i.e., log2(WT length / aberrant length). The plots are separated according to whether the change involved the creation of a splicing-derived neoepitope only (blue), the removal of a splicing-affected self-epitope only (green), or both (red). We plot in the lower panels the proportion of the total corresponding to each case.

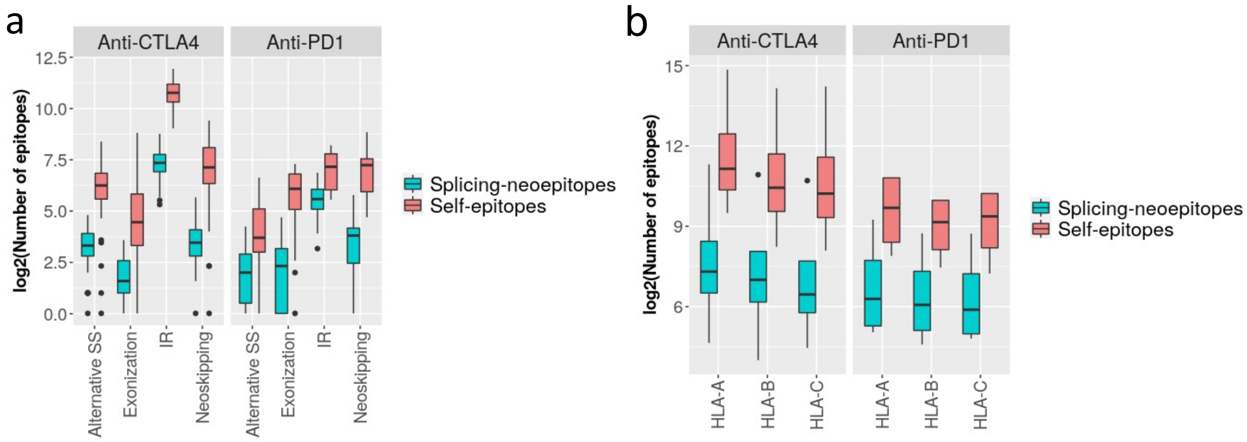


**Supp. Figure 7. Splicing-associated epitopes identified using ≤300nM**. **(a)** Distribution of the number of candidate tumor-specific splicing-derived neoepitopes (splicing-epitopes) and splicing-affected self-epitopes that would be depleted in the altered isoform (self-epitopes) using ≤300nM to define candidate epitopes. **(b)** Distribution of the number of candidate epitopes from (b), separated by HLA-type.

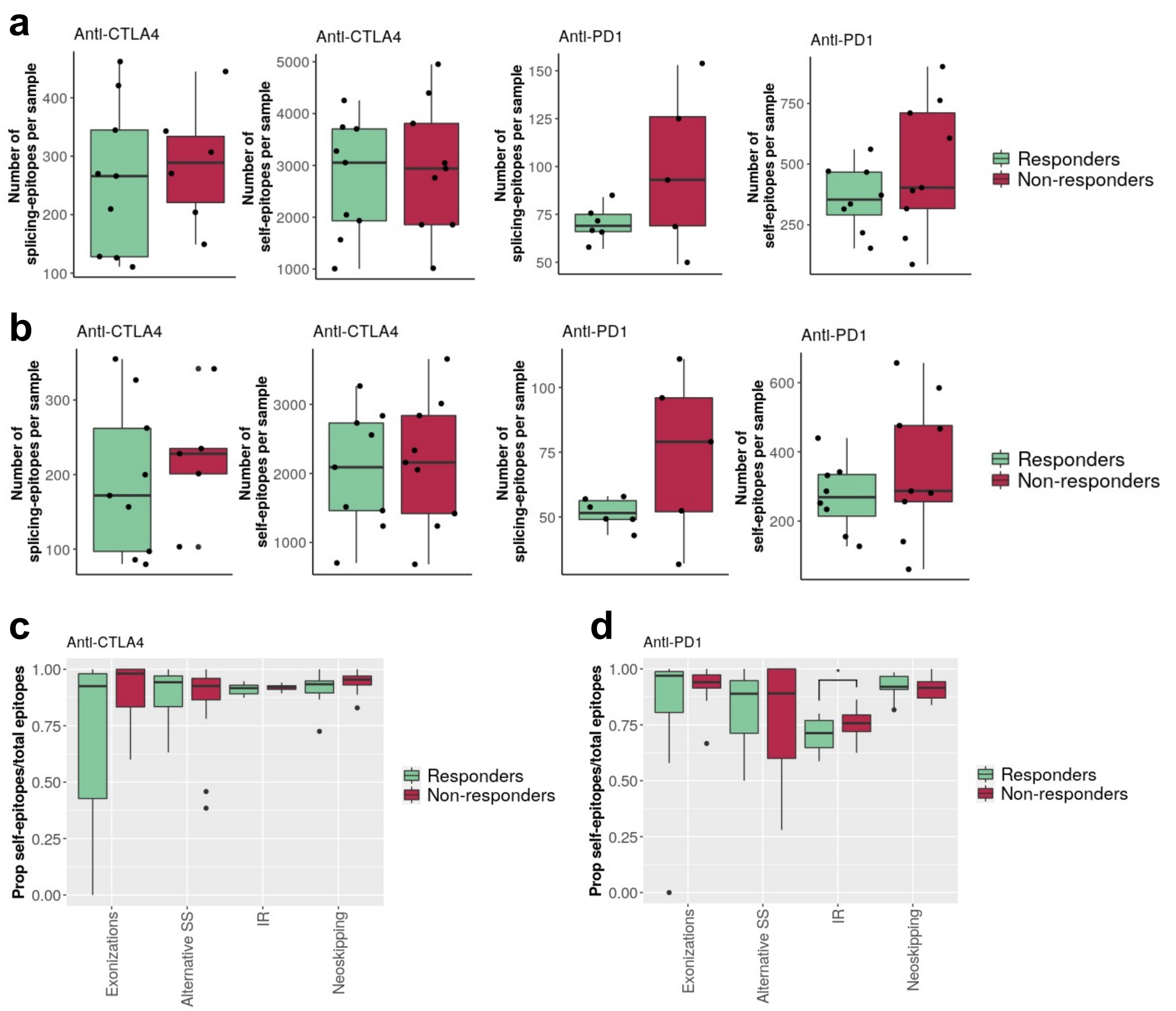


**Supp. Figure 8. Splicing-associated epitopes and immune therapy response. (a)** Distribution of the number of tumor-specific splicing-derived neoepitopes (splicing-epitopes) and splicing-affected self-epitopes that would be depleted in the altered isoform (self-epitopes) using ≤500nM to define candidate epitopes, separated by clinical outcome. The number of splicing-derived neoepitopes in responders to anti-CTLA4 (mean 346) and non-responders (mean 375) were not significantly different. Similarly, the anti-PD1 cohort showed no significant difference between the total number of splicing-derived neoepitopes between responders (mean 46) and non-responders (mean 62.6) in the anti-PD1 cohort. **(b)** as in (a) but using ≤300nM to define candidates. The number of tumor-specific splicing-derived epitopes in responders to anti-CTLA4 (median 172) and non-responders (median 228) were not significantly different. A similar result was found for the self-epitopes (2090 and 2159). We found the same for the anti-PD1 cohort (splicing tumor-epitopes: 51.5 vs 79; splicing self-epitopes: 269 vs 287). **(c)** Proportion of splicing-affected self-epitopes over the total of epitopes (splicing-affected self-epitopes and tumor-specific splicing-derived neoepitopes) (y axis) for patients treated with anti-*CTLA4*, separated by type of splicing alteration (x axis) and by patient response: responder (green) or non-responder (red). In this plot, candidate epitopes were defined using ≤300nM as threshold. **(d)** As in (b) but for melanoma patients treated with anti-*PD1*.

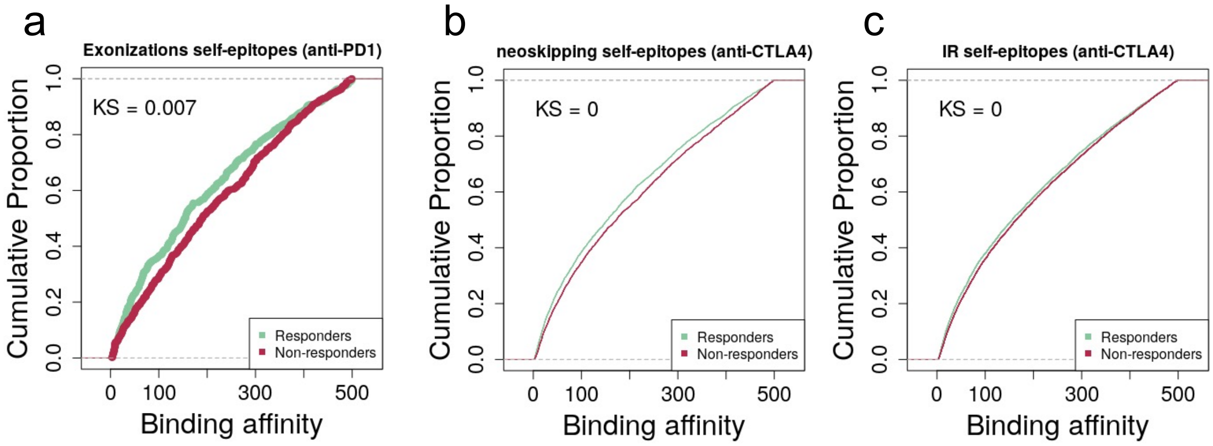


**Supp. Figure 9. Analysis of the epitope affinities in responders and non-responders using ≤500nM to define candidates. (a)** Cumulative plot of the binding affinities (x axis) of splicing-affected self-epitopes in melanoma tumors from exonization events separated in responders (green) and non-responders (red) to anti-PD1 therapy. Smaller values of binding affinity correspond to a stronger interaction between the peptides and the MHC-I complex. We also give the Kolmogorov-Smirnov test p-value (KS) = 0.0074. **(b)** Cumulative plot of the binding affinities (x axis) of splicing-affected self-epitopes in melanoma tumors from new skipping events (neoskipping) events separated in responders (green) and non-responders (red) to anti-*CTLA4* therapy, KS = 0. **(c)** Cumulative plots of the affinities of splicing-affected self-epitopes in melanoma tumors from intron retention events separated in responders (green) and non-responders (red) to anti-*CTLA4* therapy, KS = 0.

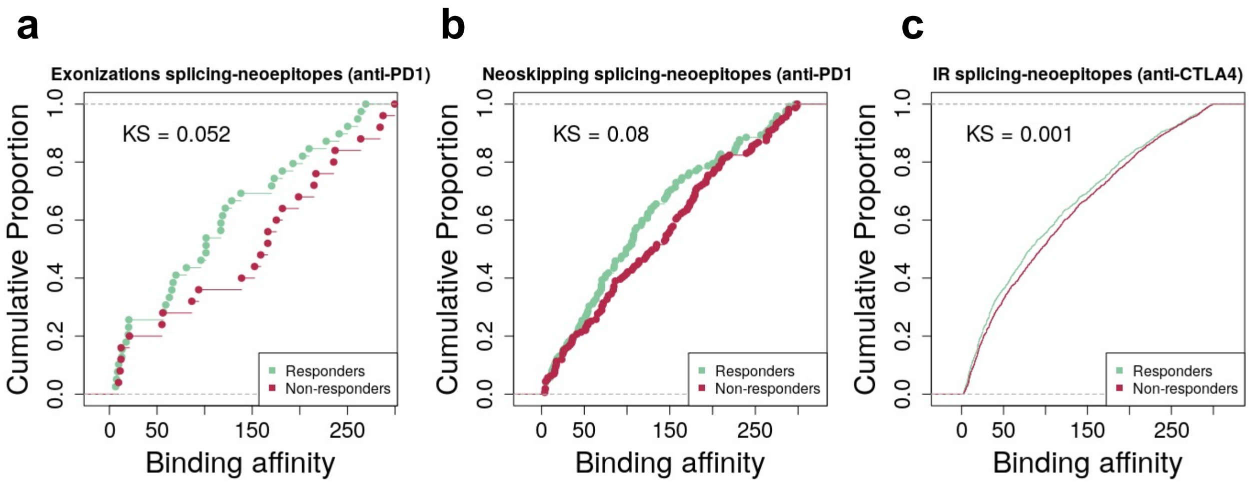


**Supp. Figure 10. Analysis of the epitope affinities in responders and non-responders using ≤300nM to define candidates. (a)** Cumulative plot of the binding affinities (x axis) of exonization-derived neoepitopes in melanoma tumors separated in responders (green) and non-responders (red) to anti-*PD1* therapy. Smaller values of binding affinity correspond to a stronger interaction between the peptides and the MHC-I complex. We also give the Kolmogorov-Smirnov test p-value (KS). **(b)** Cumulative plot of the binding affinities (x axis) of neoskipping-derived neoepitopes in melanoma tumors from separated in responders (green) and non-responders (red) to anti-*PD1* therapy. **(c)** Cumulative plots of the affinities of intron-retention-derived neoepitopes in melanoma tumors separated in responders (green) and non-responders (red) to anti-*CTLA4* therapy.

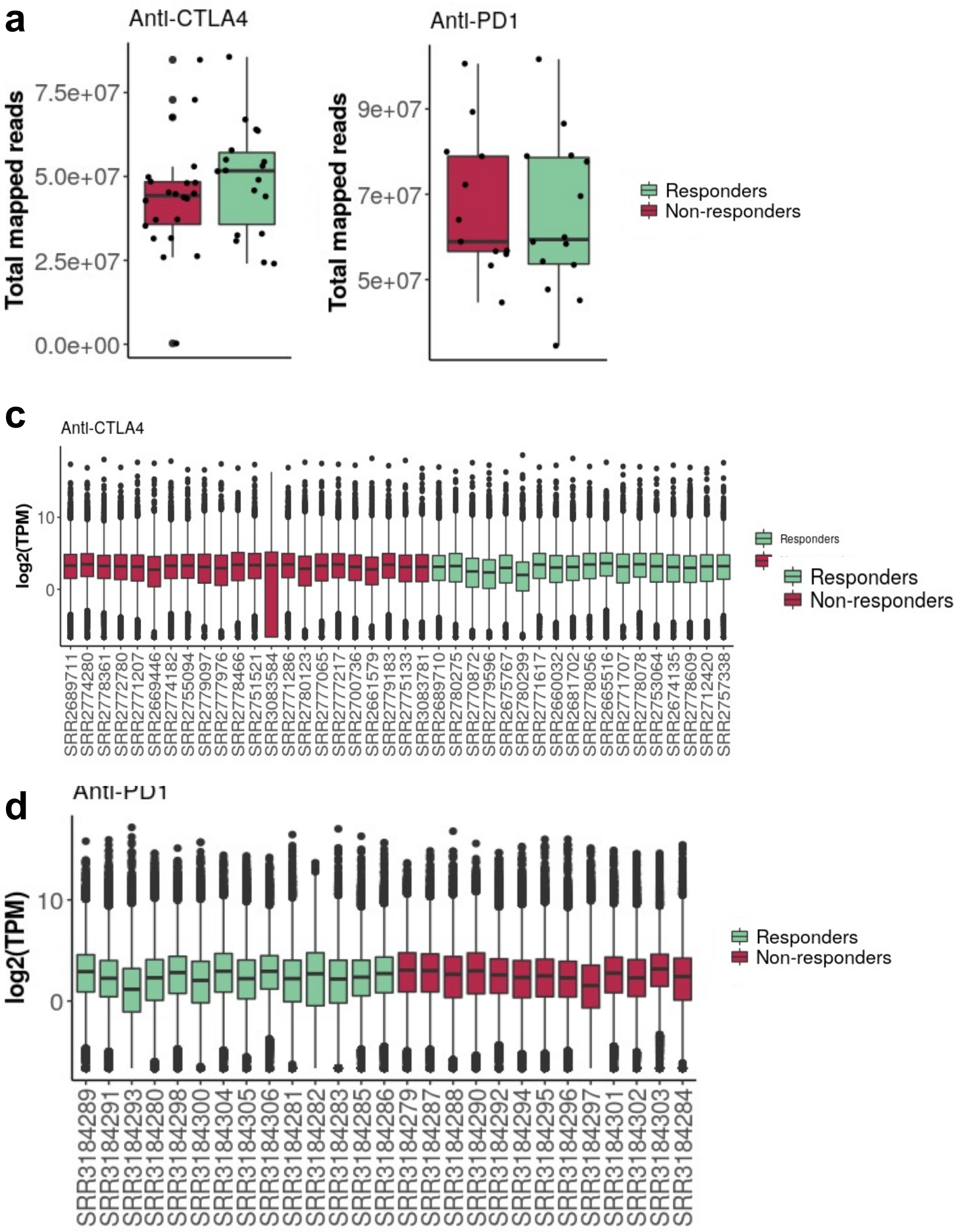


**Supp. Figure 11. Comparison of sample properties between responders and non-responders.** We show the number of mapped reads in responder and non-responder patients in the anti-CTLA4 **(a)** and the anti-PD1 **(b)** cohorts. We also show the distribution of the transcript expression values for each patient, represented as log2(TPM) (y axis) for the anti-*CTLA4* **(c)** and for the anti-*PD1* **(d)** cohorts.

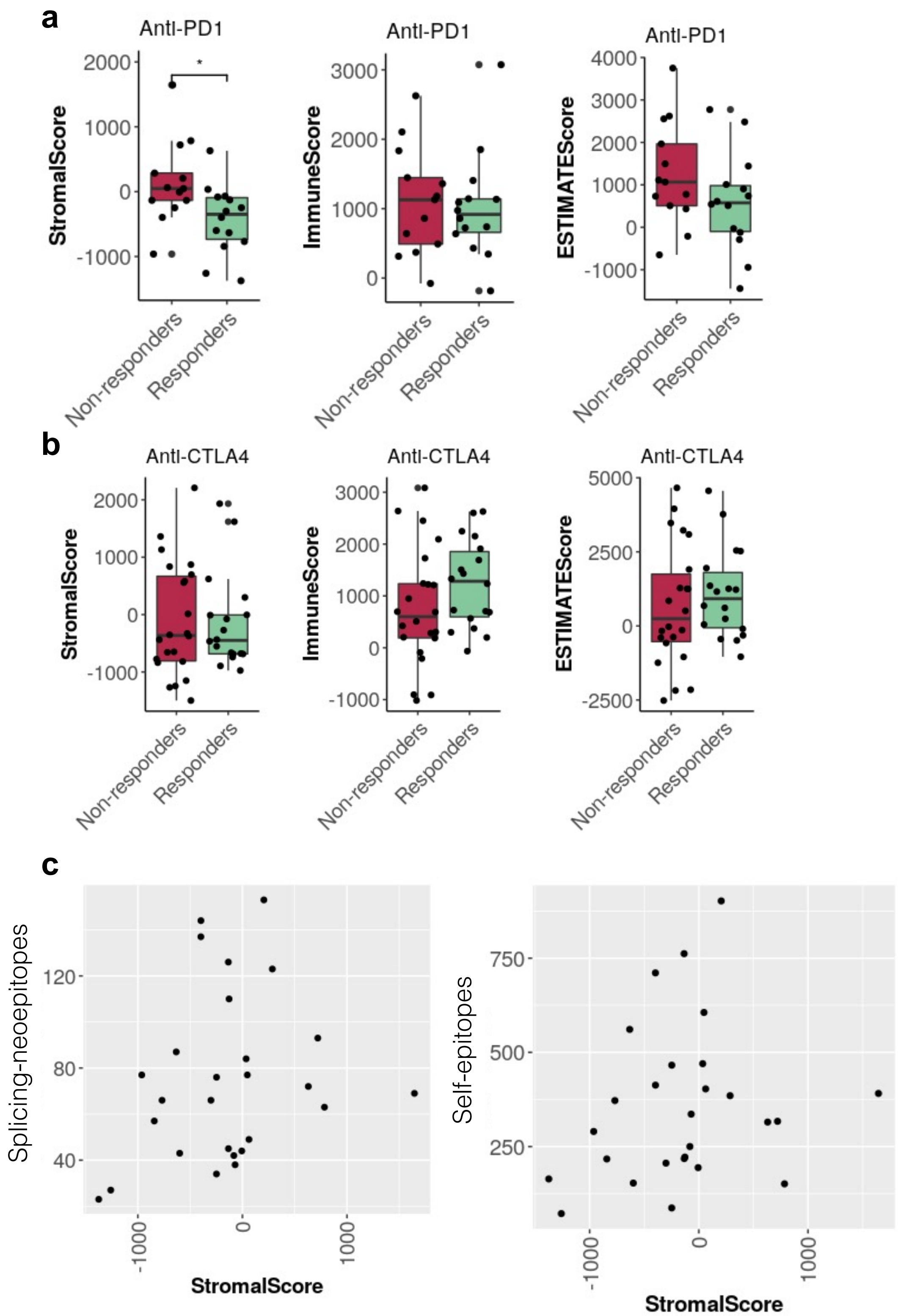


**Supp. Figure 12. Stroma and Immune content comparisons between responders and non-responders.** We show the stromal content (StromalScore), immune cell infiltration (ImmuneScore), and overall score predicted with ESTIMATE (Yoshihara et al. 2013) separating patients according to the treatment response in each cohort, anti-*PD1* **(a)** and anti-*CTLA4* **(b)**. The only significant differences detected was in relation to the stromal content in the anti-PD1 cohort (p-value ~ 0.05). **(c)**
